# Biomolecular condensates in fungi are tuned to function at specific temperatures

**DOI:** 10.1101/2023.11.27.568884

**Authors:** Benjamin M. Stormo, Grace A. McLaughlin, Logan K. Frederick, Ameya P. Jalihal, Sierra J Cole, Ian Seim, Fred S. Dietrich, Amy S. Gladfelter

## Abstract

Temperature can impact every reaction and molecular interaction essential to a cell. For organisms that cannot regulate their own temperature, a major challenge is how to adapt to temperatures that fluctuate unpredictability and on variable timescales. Biomolecular condensation offers a possible mechanism for encoding temperature-responsiveness and robustness into cell biochemistry and organization. To explore this idea, we examined temperature adaptation in a filamentous-growing fungus called *Ashbya gossypii* that engages biomolecular condensates containing the RNA-binding protein Whi3 to regulate mitosis and morphogenesis. We collected wild isolates of *Ashbya* that originate in different climates and found that mitotic asynchrony and polarized growth, which are known to be controlled by the condensation of Whi3, are temperature sensitive. Sequence analysis in the wild strains revealed changes to specific domains within Whi3 known to be important in condensate formation. Using an *in vitro* condensate reconstitution assay we found that temperature impacts the relative abundance of protein to RNA within condensates and that this directly impacts the material properties of the droplets. Finally, we found that exchanging Whi3 genes between warm and cold isolates was sufficient to rescue some, but not all, condensate-related phenotypes. Together these data demonstrate that material properties of Whi3 condensates are temperature sensitive, that these properties are important for function, and that sequence optimizes properties for a given climate.

## Introduction

Temperature can impact every reaction that is essential to the life of a cell. The structure of proteins and nucleic acids^1^, the fluidity of membranes^2^ and the rate of diffusion^3,4^ are all temperature-dependent processes. For sessile and ectothermic organisms that cannot regulate their own temperature, a major challenge is how to ensure cellular function is robust to temperatures changes that are unpredictable and occur on multiple timescales. While there are specific examples of molecular temperature sensors such as RNA thermometers^5^, heat shock proteins^6,7^, and the unfolded protein response^8^, how cells sense and adapt to temperature changes happening across broad ranges of temperatures and on varying timescales remains poorly understood. Phase transitions involving biomolecules condensing in cells offer a possible general mechanism for encoding temperature-responsiveness into cell biochemistry and organization.

Biomolecular condensates are often driven to form by multivalent interactions amongst proteins with intrinsically disordered regions (IDR) and nucleic acids^9,10^. The formation of and resulting properties of condensates are dependent on both the valency and strength of interactions which in turn are dependent on the temperature of the system. In several systems including the *Drosophila* nucleolus^11^ and P-granules in *C. elegans*^12^ the formation of condensates has been shown to depend on the temperature of the system. Condensates have also been implicated as temperature sensors able to respond to changes in temperature by phase separating. It has been shown in *Arabidopsis* that the poly-Q containing protein ELF3 varies in sequence in accordance with climate allowing for thermal control of ELF3 condensation^13^. It has also been shown that the yeast protein Ded1p undergoes a phase transition as the cells approach their maximal growth temperature and that this phase transition varies between diverse yeasts with different temperature preferences ^14^. Similarly, the IDR sequences of the C-terminal domains of RNA polymerase II from cold adapted yeasts is associated with the ability to phase separate at different temperatures^15^. These examples suggest that temperature-responsiveness can be sequence-encoded in IDRs that form biomolecular condensates in sessile organisms.

Intriguingly, simple eukaryotes, which are more likely to be ectothermic, have a larger percentage of disorder in their proteomes compared to more complex eukaryotes. Further, parasites that host-switch, experiencing both ambient and host body temperatures, have the highest percentage of IDR-containing proteins, supporting the hypothesis that these types of sequences may be critical to adapt to rapidly changing environments^16^. The presence of disordered regions is however only one way to promote multivalent interactions that drive condensate formation. Another critical avenue is through protein-nucleic acid binding. Many condensates contain nucleic acids, and IDR-containing nucleic acid binding proteins often bind to a specific motif or structures in nucleic acids^17^. These sequence-encoded interaction sites can potentially be a valuable substrate for evolution because mRNA sequence can evolve independently of the protein they code for^18,19^. Similarly, IDRs may be less evolutionarily constrained than folded proteins because of their ability to sample many conformations and not be restricted to one functional shape^20,21^. The properties of proteins and nucleic acids that participate in biomolecular condensates make them prime candidates as vehicles for adaptation of sessile organisms to fluctuating environmental conditions.

Filamentous fungi are a major component of the terrestrial biosphere and integral to biogeochemical cycles, underlying the importance of understanding how these organisms respond to fluctuating environments^22,23^. *Ashbya gossypii* is a fungus that is closely related to *Saccharomyces cerevisiae* but grows exclusively by filamentous growth as a syncytial mycelium instead of as a uninucleate yeast^24^. Within the hyphal cytoplasm are numerous nuclei which undergo mitosis independently of one another and multiple separate points of polarized growth^25,26^. *Ashbya’s* continuous cytoplasm is thus compartmentalized with respect to growth and nuclear division. Key to this compartmentalization is the protein Whi3. Whi3 forms condensates throughout the cytoplasm of *Ashbya* in the vicinity of nuclei and at sites of polarized growth^27–29^ (**FIG 1A**). In *Ashbya,* at least three distinct Whi3 condensates are known and are characterized by the RNA molecules in the condensates and the functions of the condensate. Whi3p complexes with the RNAs *SPA2* and *BNI1,* a polarity scaffold and a formin, near hyphal tips and near nascent branch sites. These complexes are important for maintaining tip growth and for generating new lateral branches. Loss or mutation of Whi3 leads to *Ashbya* with unbranched hyphae^27^. Whi3 also forms distinct foci with the cyclin RNA *CLN3* in *Ashbya* and *S. cerevisiae*. In both species these complexes have been shown to position *CLN3* near nuclei^28,30^. In *Ashbya* these *CLN3* condensates are required for asynchronous mitosis such that mitosis becomes synchronous in *whi3* mutants or in *cln3* mutants in which the Whi3 binding site has been mutated. These different condensates have varied properties *in vitro* and *in vivo* that contribute to their distinct molecular compositions despite all sharing the Whi3 protein^31,32^.

**Figure 1:**
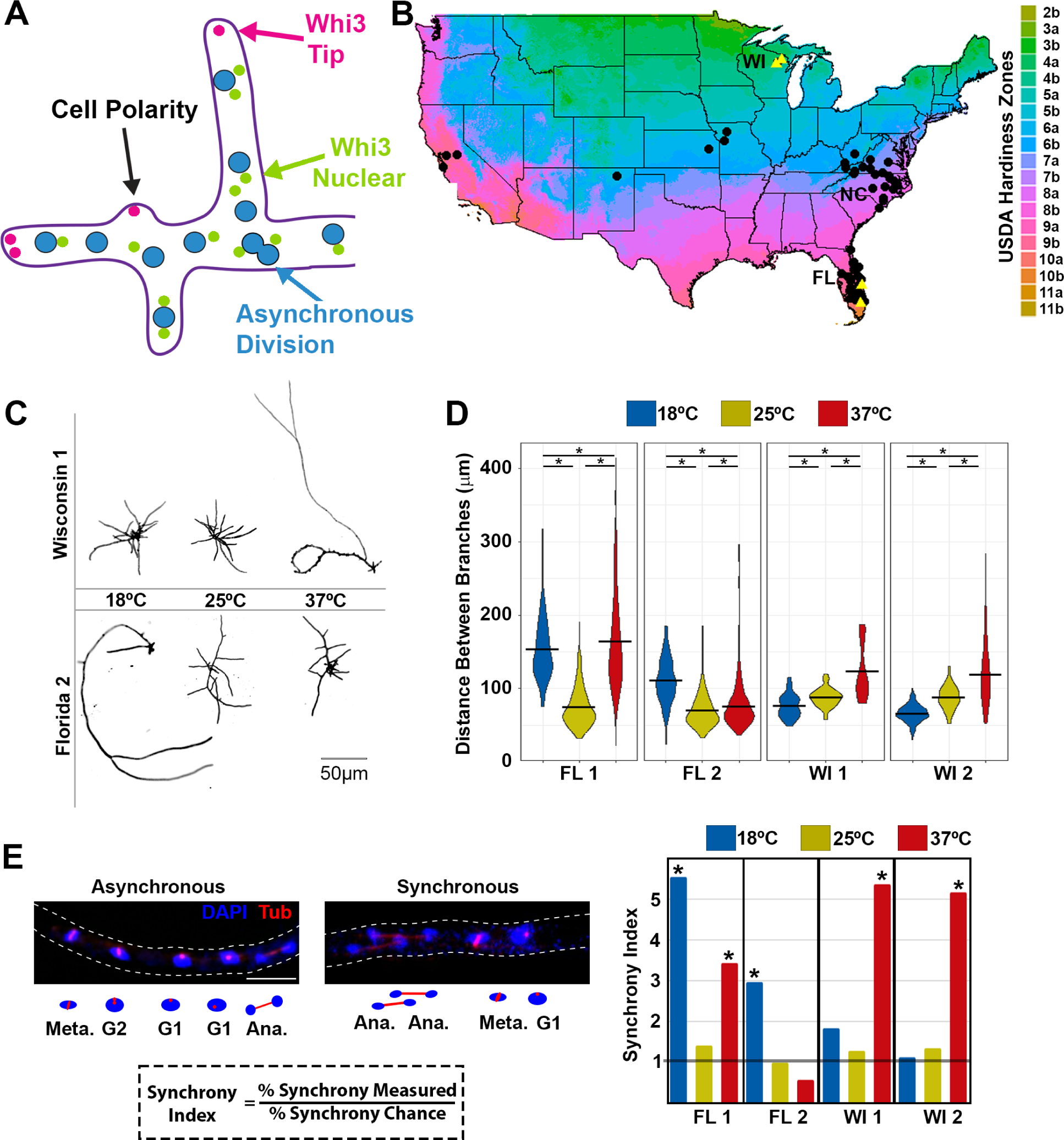
Temperature-sensitive phenotypes in *Ashbya* isolates from different climates. **A)** schematic of an *Ashbya* cell showing nuclei (blue) and Whi3 nuclear-associated condensates (green) and polarity-related condensates (magenta) all within a single hyphae (purple outline). An asynchronous mitosis is shown. **B)** location of wild isolates from around the United States (black dots) with the isolates chosen for further phenotypic characterization in yellow triangles. Points are jittered to avoid overlap and so do not correspond to exact locations. Map colors represent USDA hardiness zones ^40^ **C)** Representative images of hyphae stained with a cell wall stain calcofluor white from the indicated isolates grown at low, medium and high temperatures. **D)** Quantitation of the distance between branches was measured and an average branch length for each cell was calculated for each isolate grown at the indicated temperature. (ANOVA * P<0.05) **E)** Representative immunofluorescence of the spindle poll body (SPB) to score mitotic synchrony. Tubulin (red) and DAPI (blue) from an asynchronous hyphae and a synchronous hyphae, the outline of the hyphae is indicated by the dashed lines. Below is a schematic of the same hyphae with cell cycle state indicated for each nucleus. The synchrony index is the ratio of actual number of paired mitotic nuclei divided by the number expected by chance, which is the square of the total proportion of nuclei in mitosis. Quantitation of the synchrony index from each isolate at the indicated temperature, the line at 1 indicates random. Scale bar indicates 5µm. (t-test * is P<0.05).

Ashbya is found as a free-living fungus or in contact with insects in the genera *Oncopeltus,* which are thought to be purely ectothermic as well^33,34^. Thus, *Ashbya* experiences ambient temperature fluctuations during its life cycle. We predicted that based on the wide-geographical ranges of *Ashbya* and its insect host and the critical role that biomolecular condensates play in the growth of this fungus that this was a good candidate for examining the role of condensates in temperature adaptation. Previous work with Whi3 condensates reconstituted *in vitro* indicate the assemblies are formed by three main types of interactions: 1) Whi3-Whi3 protein interactions which depend on a Glutamine-Rich Region (QRR), 2) Whi3-RNA interactions take place through an RNA Recognition Motif (RRM) in Whi3 which binds the motif UGCAU in target RNAs and 3) RNA-RNA interactions which take place due to base-pairing between RNAs ^27,31,32,35^. These homotypic and heterotypic interactions offer a broad range of sequence space where changes could modify interaction strengths and temperature sensitivity of Whi3 condensates. In this study we examine temperature sensitivity of *Ashbya* derived from different climates and find evidence supporting that sequence-encoded changes in Whi3 enables adaptation to growth at varying temperatures.

## Results

To begin investigating the relationship between temperature sensitivity and *Ashbya* cell growth, we assessed a collection of 70 *Ashbya* strains from different climate zones within the United States containing both previously studied isolates and new isolates^33^ (**FIG 1B**). We focused in-depth analysis on a subset of these isolates, reasoning that adaptation to specific temperature regimes would be most pronounced in isolates from the most different climates (Wisconsin, latitude 43° N, which has colder winters and milder summers, abbreviated WI and Florida, latitude 25° N, which has a warm subtropical climate year-round, abbreviated FL) (**FIG 1B** yellow triangles).

### Geographical origin associated with distinct temperature-sensitive phenotypes

We first examined if cell branching and mitotic synchrony, which both require Whi3 condensates, varied between isolates from different climates when grown at different temperatures. Previous work showed that *Ashbya* lacking Whi3 protein entirely, lacking the QRR domain, or with point mutations in the RRM that prevent RNA binding all have defects in cell polarity in which mycelia contain long hyphae with few lateral branches ^27^. We grew a subset of FL and WI isolates at low (18°C), moderate (25°C), and high (37°C) temperature and then measured the distance between branches along the mycelia (typical lab strain culturing is performed at 30°C). We found that isolates from cold climates (WI 1 and WI 2) had temperature sensitive polarity problems marked by large interbranch distances at hot temperatures compared to cold temperatures (**FIG 1C, D**). Conversely, both warm isolates (FL 1 and FL 2) had large interbranch distance at low growth temperatures. One warm isolate (Fl 1) exhibited increased inter-branch distance at both high and low temperatures (**FIG 1 C, D**). These results suggest that branching morphology in *Ashbya* cells is temperature sensitive and that this sensitivity is associated with the climate from which the isolate originated.

We next assessed if mitotic synchrony was temperature sensitive in wild strains. To measure nuclear synchrony, we performed immunofluorescence against tubulin to detect the spindle pole body (SPB) which changes in characteristic ways throughout the cell cycle^25,36^ (**FIG 1E)**. We categorized each nucleus as having one SPB (G1), duplicated SPB (S/G2), or a spindle (M-phase) and then measured the proportion of nuclei in each class. By squaring the proportion of nuclei in M-phase we could calculate the probability that two neighboring nuclei would both be in mitosis by chance. We compared this to the actual number of paired mitotic nuclei to calculate the “synchrony index” which is the fold enrichment for mitotic pairs^25^. A synchrony index of one indicates that neighboring nuclei are in a random cell cycle position relative to their neighbors, and this is the typical state for the reference strain at 30°C^25,37–39^. Higher synchrony index numbers indicate synchrony between neighbors above the level expected by chance. All four examined isolates exhibited asynchronous mitosis at moderate (25°C) temperatures in agreement with previous studies on the reference strain^25,28^. We found that the cold climate isolates (WI 1 and WI 2) were also asynchronous when grown at low temperatures (18°C) but became synchronous at high temperatures (37°C). Conversely, one of the warm isolates (FL 2) was asynchronous at warm temperatures but displayed increased synchrony at low temperatures. As with the branching phenotype, the other isolate from Florida (FL 1) was defective at both high and low temperatures (**FIG 1E**).

Thus, temperature sensitive phenotypes for polarity and mitosis were seen at both high and low temperatures in an isolate-specific manner. Interestingly one isolate (FL 1) displayed both cold sensitivity and heat sensitivity. The behavior of this isolate suggests that the temperature sensitive phenotypes can arise in cells that are too far above or below the optimal range, but the width of the specific optimum range has some variability even within a climate zone. Together these data suggest that Whi3-dependent cell processes fail to function properly outside of a narrow range of temperatures and instead are similar in phenotype to a *WHI3* loss of function at temperatures outside this optimum.

### Isolates vary in Whi3 Q-tract length, codon use and binding sites in target RNAs

The cell morphology and cell cycle phenotypes are suggestive of temperature sensitivity in Whi3-dependent processes, so we next looked for changes to the sequence of *WHI3, CLN3* and *BNI1* in the wild-isolates. We performed Sanger sequencing on 70 total isolates and found 11 unique Whi3 sequences (**FIG 2 A,B)**. Alignments of these sequences revealed substantial differences between isolates. Notably, the majority of changes to the Whi3 protein sequence were located within the QRR **(FIG 2A, Fig S1A),** with the number of glutamines varying from 111 to 128 (**FIG 2B, FIG S1A,D**). Based on previous work in *Arabidopsis* on ELF3, we had initially predicted that isolates from warmer climates (Florida) would have more glutamines in their QRR ^13^. However, we found that the selected WI isolates from colder climates contained 120 and 124 glutamines compared to 112 and 113 glutamines in the selected Florida isolates. Interestingly, the expanded QRR in colder isolates is associated with the loss of a heptad repeat sequence that we have previously shown to oppose phase separation through an oligomerization mechanism ^35^. The isolates containing more glutamine residues contain a mutated heptad repeat lacking one of the three leucines needed to form a coiled-coil domain (**FIG S1B,C,E)**. Notably, we also found *WHI3* genes that code for different numbers of glutamines can exist in close geographic proximity, for example, both the maximum and minimum number of glutamines were from the same state (North Carolina, abbreviated NC) however these examples were from different geographical and climate areas of the state. The isolate containing fewer Qs was found in the eastern more temperate coastal area and the greater number of Qs found in the mountainous western area, a different and colder climate zone. The large variability within NC also likely results, in part, from sampling bias, because almost one third of the isolates are from that state. Together these data show there are substantial changes in the Whi3 protein, particularly in the QRR, amongst isolates from different climates that are consistent with the temperature-sensitive phenotypes for Whi3-dependent processes.

**Figure 2:**
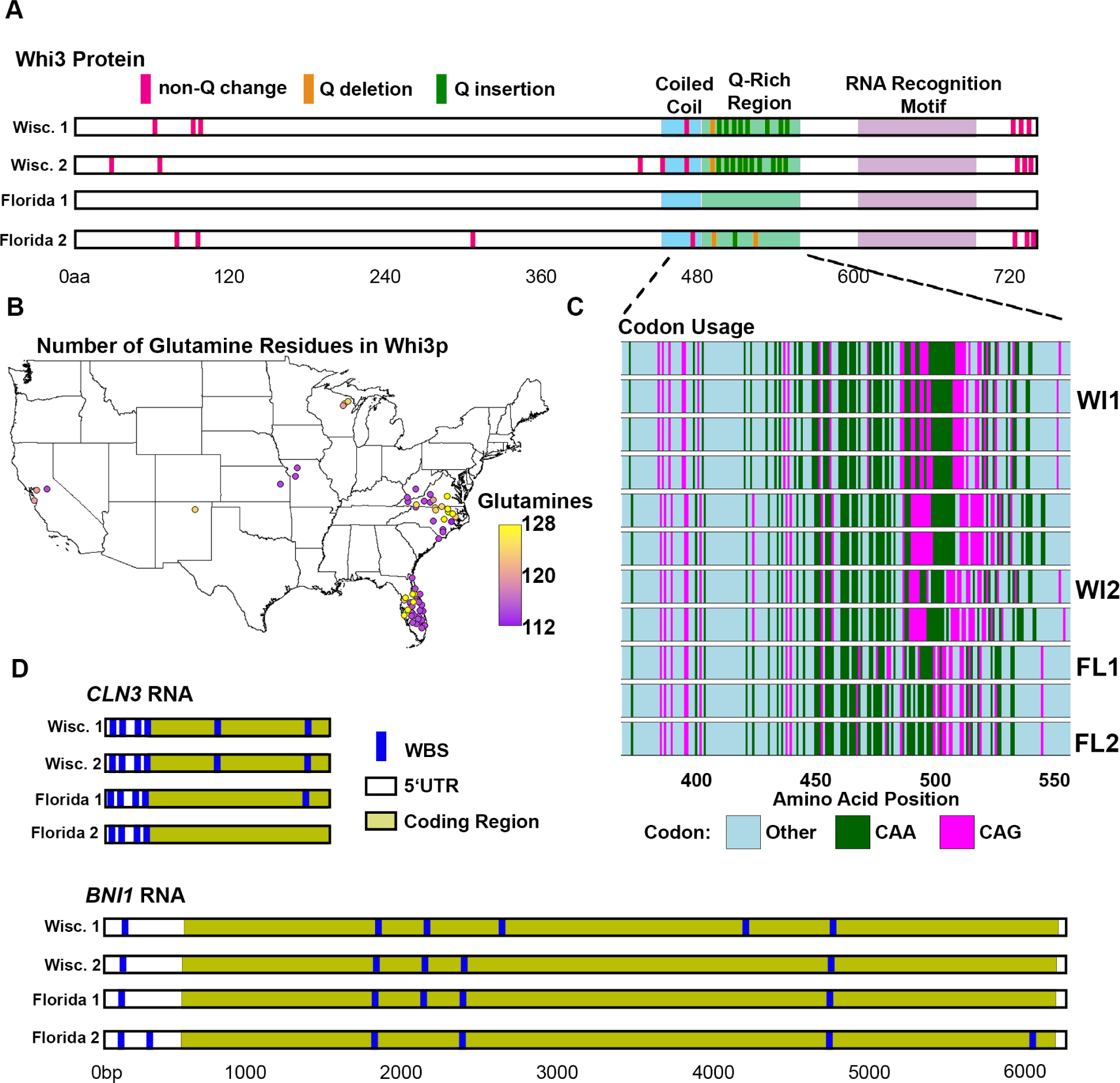
*Ashbya* isolates show sequence changes in components involved in Whi3 condensate formation. **A)** Alignment of Whi3 sequence showing the coiled-coil (light blue), Q-Rich Region (green) and the RNA Recognition Motif (purple). Protein coding changes between the isolates are shown relative to the reference strain with non-glutamine substitutions (pink), glutamine insertions (green), and loss of glutamine (orange) noted. Most of the changes are confined to the QRR. Numbers indicate amino acids position. **B)** Map of the United States showing the location of all sequenced Whi3 color coded by the number of glutamines present in the entire protein. **C)** Schematic of the QRR showing the codon usage for glutamine CAA (green) and CAG (magenta). Both codons are used throughout the protein and the glutamines are separated by non-glutamine amino acids. Numbers indicate amino acid number from the start of the protein. The four strains used are indicated. **D)** Schematic alignments of *BNI1* and *CLN3* RNAs showing the coding region (yellow) the predicted UTR (white) and the Whi3 binding motifs (UGCAU). Numbers indicate the number of nucleotides from the predicted start of transcription.

The coexistence of strains with different *WHI3* sequences in geographic proximity suggests that there is either genetic instability in the QRR, some genetic exchange across the different sampling areas and/or other avenues for adapting Whi3 condensate functions apart from changes to *WHI3*. If the QRR is highly labile and changing quickly, we would predict that there might be substantial tracts of glutamines that are coded for with the same codon as such repetitive sequences are known to be unstable ^41^. However, we find that *WHI3* has both CAG and CAA codons and that these codons are intermixed in the QRR. This suggests that in fact the QRR may be somewhat stabilized (**FIG 2C, FIG S1A**). Previous work has identified several phosphorylation sites within Whi3 that have phenotypic effects when mutated, however, none of these known phosphorylation sites differed between the wild isolates ^39^(data not shown). Thus, the sequences of Whi3 vary substantially across the population of wild-isolates and much of the variation is harbored in the QRR, with longer QRRs associated with the absence of an oligomerization domain.

We next aligned the sequences of the RNAs that are known to phase separate in complex with Whi3 protein. We focused on the Whi3 binding motifs in *CLN3* and *BNI1*, noticing changes to both the number and position of Whi3 binding motifs within these RNAs (**FIG 2D**). We predicted that warmer isolates would have more binding sites, based on the idea that higher temperature condensation might require higher valence amongst molecules, however we found that for *CLN3* the warmer isolates had 4 or 5 binding sites whereas both Wisconsin isolates had 6. For the *BNI1* transcript we found a more ambiguous results with one Florida and one Wisconsin isolate containing 6 binding sites and the other Florida and Wisconsin containing 5 binding sites although these were not the same sites (**FIG 2D**) ^32^. These data suggest that the components of Whi3 condensates accumulate changes that can alter valency and likely the strength of interactions amongst key components that could drive the temperature-sensitive functions of Whi3 condensates.

### Condensates in cells vary in frequency and position as a function of temperature

Given the sequence changes and striking temperature-sensitive phenotypes in Whi3-dependent processes, we next sought to observe the condensates within cells. To do this we tagged Whi3 in a warm and a cold isolate (WI 1, FL 2) at the endogenous locus using the fluorescent monomeric protein mNeon to generate strains we called WI^Whi3-mNeon^ and FL^Whi3-mNeon^ ^42^. mNeon was chosen because of its combination of brightness and because it does not exhibit a propensity to dimerize, which can affect the behaviors of phase separating proteins ^43^.

We were first interested in whether the total amount of Whi3 protein changed at different temperatures since condensate assembly may be sensitive to the concentration of critical components. Conceivably changes in protein levels could serve as a rapid and sequence-independent way for these cells to adapt to different temperature changes. We measured the total fluorescence within the hyphae to estimate protein level and found that protein levels were relatively constant across temperatures (**FIG 3A,B**) eliminating this as a likely mechanism explaining the temperature sensitivity.

**Figure 3:**
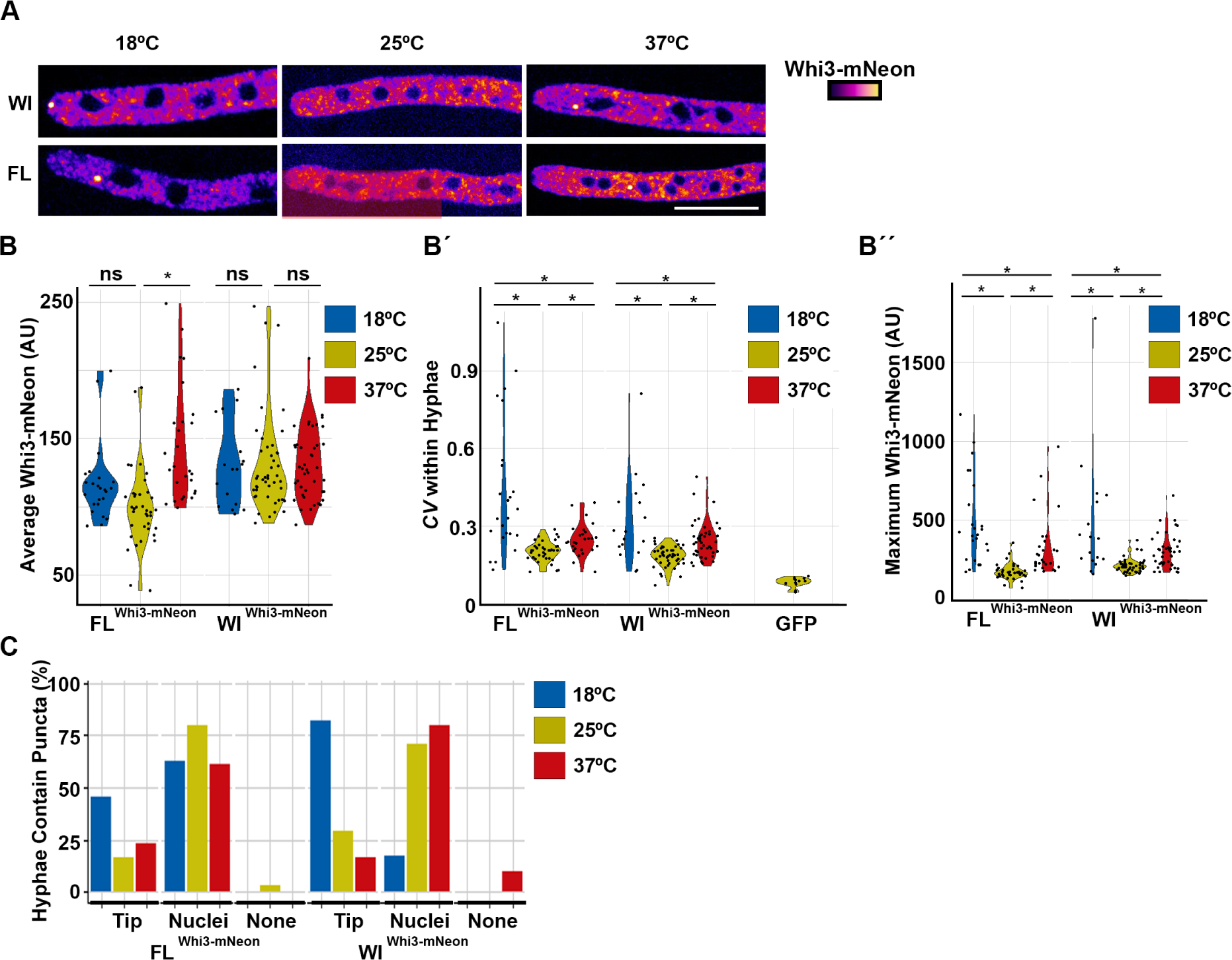
Condensates vary between isolates and across temperatures *in vivo.* **A)** Representative images of hyphae from both isolates at each temperature. Whi3-mNeon signal intensity is fire LUT coded. Scale bar represents 10µm. **B)** Quantification of Whi3 protein concentration for each isolate across temperatures was determined by measuring the fluorescence intensity of mNeon within the hyphae. **B’)** Coefficient of Variation (*CV)* for fluorescence intensity along the hyphae in each indicated strain at all three temperatures along with a cytoplasmic GFP control. **B’’)** The maximum Whi3 intensity was determined by measuring fluorescence within the hyphae **C)** Quantification of the percentage of hyphae that contain condensates near tips and near nuclei or that lack condensates (none) for each isolate at different temperatures.

We next analyzed the appearance of condensates at different temperatures. Whi3 was highly heterogenous with bright puncta and a dimmer but also uneven cytosolic signal **(FIG 3A,B’).** Since condensates can form with sizes both above and below the diffraction limit, we used variation in pixel intensity to estimate the degree of condensation (reported as coefficient of variation, *CV*). Because Whi3 is nuclear excluded, we only measured pixel values in the cytoplasm. As a control, we also measured the *CV* of a cytoplasmic GFP which was uniform throughout hyphae (**Fig 3B’**). In both the WI and FL isolates, we found the highest heterogeneity in signal at 18°C and the lowest heterogeneity at 25°C, which was a significantly different distribution of CVs based on a K-S test. This effect can also be seen by measuring the intensity of the brightest foci, which fall within a narrow range at 25°C but hyphae at 18°C and 37°C tend to have much brighter foci, suggesting more intense foci with more variation both within and between hyphae are associated with loss of function at sub-optimal temperatures (**Fig 3B’’)**.

To assess protein localization after an acute temperature change, we shifted cells grown at 18°C to 37°C and cells grown at 37°C to 18°C and then imaged after 10 min at the new temperature. We reasoned that such a short time frame could reveal temperature-dependent properties without a chance for the cell to alter transcription or translation. We first checked the protein level as a control because we predicted that a short shift would not affect total protein level, which we confirmed by measuring total fluorescence (**FIG S2A**). We then measured the *CV* and found a decrease in heterogeneity in cells that were shifted from cold to hot but no change in cells shifted from hot to cold (**Fig S2B)**. This suggests that in the absence of translation and transcription, Whi3 condensates are intrinsically more unstable at high temperature compared to lower temperatures as would be expected for a system with upper critical solution temperature behavior. In summary, these data show that Whi3 assemblies change at different temperatures, but the impact is a redistribution into different scaled assemblies, many of which may be below the diffraction limit, rather than a binary change in presence or absence of the assembly. Furthermore, the presence of foci containing more protein were associated with conditions with loss-of-function phenotypes.

As various Whi3 functions are dependent on differently localized Whi3 condensates (**FIG 1A**), we also assessed if there were temperature-dependent changes in Whi3 subcellular location. We therefore categorized hyphae based on whether bright condensates were localized at the tip, near nuclei, or absent (**FIG 3C).** At high temperatures, we found a decrease in tip-localized puncta and an increase in hyphal puncta, which are commonly near nuclei. This trend was most clear in the cold climate WI isolate. These trends suggest that larger Whi3 puncta around nuclei may not only be associated with normal function but rather the brighter assemblies appear linked to a loss of function. This could be that at non-optimum temperatures large assemblies regulating the nuclear division cycle are losing normal functions or have gained a mis-regulated negative function. Importantly, it reveals differences in the temperature-sensitive properties of the tip and nuclear-associated puncta and that the presence or absence of a large condensate cannot be equated to activity that promotes asynchrony and branching. Together these data point to the specific size and properties of Whi3 condensates being critical for function, not simply the presence or absence.

### Size and protein content of condensates varies with temperature *in vitro*

The sequence differences in the QRR region of Whi3 protein and binding site valence in target RNAs as well as the condensates seen *in vivo* indicate that temperature sensitivity could be intrinsic to the Whi3 protein and relevant RNA interactors. To address this possibility, we assessed the ability of proteins and RNAs from the different isolates to phase separate at different temperatures *in vitro* using a reconstitution system previously developed by the lab ^31^. To do this we recombinantly expressed the Whi3 proteins and *in vitro* synthesized RNA for the predicted transcripts of *BNI1* and *CLN3* from the Wisconsin isolate (WI 1) and the Florida isolate (FL 2) (**FIG S3A**,**B)**. We then mixed protein with RNA from the same geographical origin and incubated at high (37°C), moderate (25°C) or low (18°C) temperature and imaged the resulting condensates **(FIG 4 A,C**). By measuring the fluorescence intensity in the center of the resulting condensates, we could compare the relative abundance of Whi3 protein and *CLN3* in the dense phase, recognizing that there may be difficulty in interpreting the absolute concentration due to potential quenching of fluorophores. We found that temperature had a dramatic effect on the resulting condensates. For condensates formed with *CLN3* RNA, the ratio of protein:RNA increased as temperature increased for both isolates (**FIG 4 B,B’).** This change was mostly due to a decrease in the concentration of *CLN3* within the droplets. While this trend was consistent between the isolates, there were substantial differences in the distribution of protein to RNA ratios and the maximum and minima of the populations between the different isolates. For condensates formed with *BNI1* RNA, we again found the lowest protein:RNA ratio at 18°C but the highest ratio was at 25°C (**FIG 4D,D’)**. These results indicate general temperature sensitivity of the Whi3-RNA system and show that temperature can alter the composition of these condensates.

**Figure 4:**
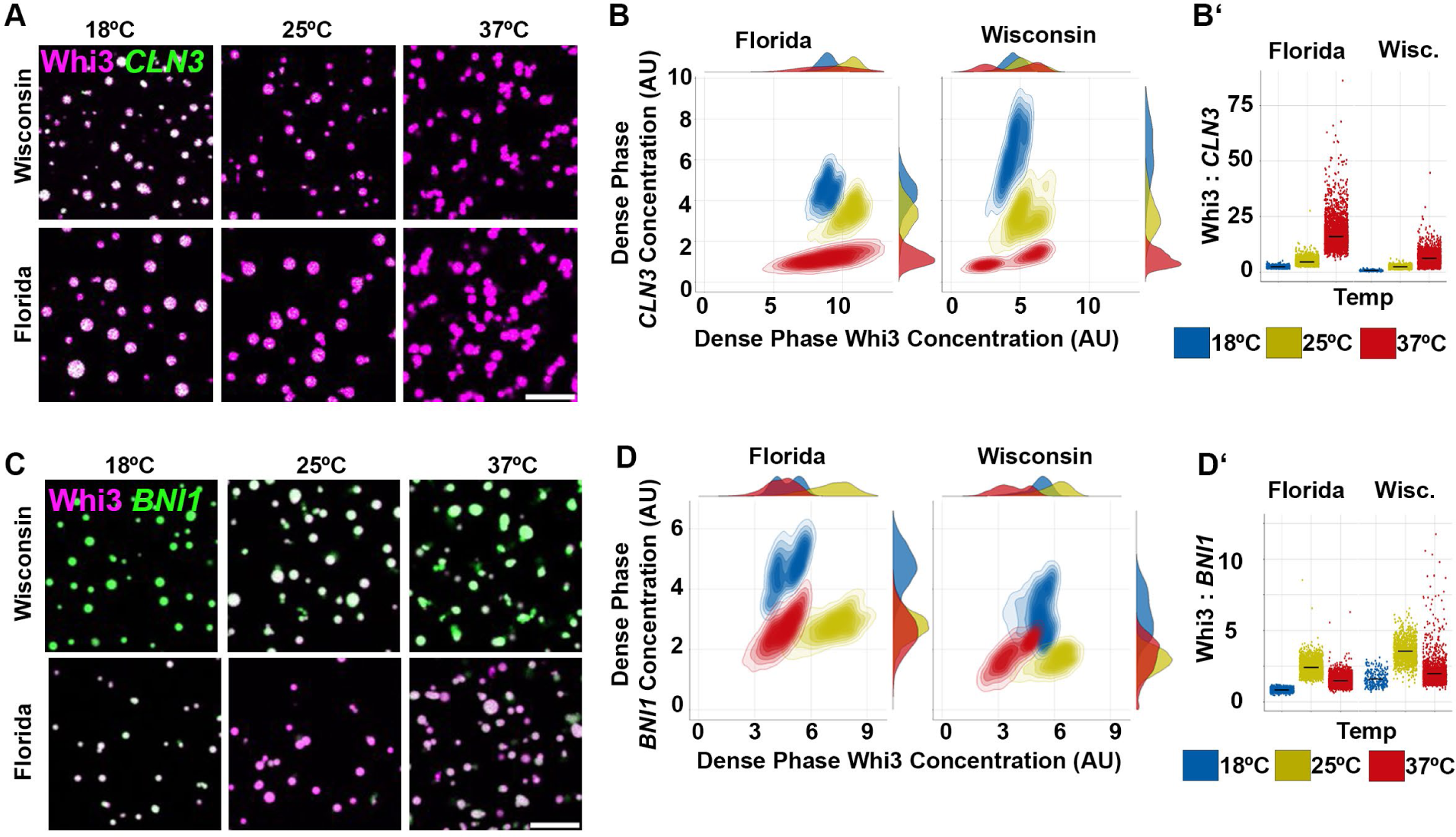
Temperature-sensitive behavior of condensates in vitro **A)** Representative images of *in vitro* condensates made with purified protein and *in vitro* transcribed RNA from the indicated isolate at each temperature. Atto-488 labelled Whi3 protein in magenta, Cy-5 UTP labelled *CLN3* is in green. **B)** Quantification of dense phase protein and RNA concentrations at each temperature from an initial bulk concentration of 1µM Whi3 and 4nM *CLN3.* **B’)** The ratio of Whi3 to *CLN3* RNA within the dense phase. Each dot represents one droplet. **C)** Representative images of *in vitro* condensates made with purified protein and *in vitro* transcribed RNA from the indicated isolate at each temperature. Atto-488 labelled Whi3 protein in magenta, Cy-5 UTP labelled *BNI1* is in green. **D)** Quantification of dense phase protein and RNA concentrations at each temperature from an initial bulk concentration of 1µM Whi3 and 4nM *BNI1*. **D’)** The ratio of Whi3 protein to *BNI1* RNA within the dense phase of droplets. Scale bars represent 5µm.

Previous work in *Ashbya* found that high protein:RNA levels were associated with more liquid-like droplets ^31,35^. To measure the material properties of droplets made from wild-isolate sequences we performed three tests. We first assessed the shape of the droplets as droplets that are more liquid-like will fuse upon contact to minimize surface tension in contrast to more viscoelastic droplets that will only slowly fuse or not fuse under the time scale of the experiment leading to non-circular assemblies^44,45^. We measured the circularity of the Whi3 wild-isolate condensates (circularity is defined 4π* Area/Perimeter^2^ so that a perfect circle equals 1) and found that as temperature increased, and protein:RNA increased, circularity actually decreased (**FIG S3D,G)**. Visually, the non-circular condensates at high temperatures are formed of chains of droplets which contacted each other and failed to relax (**FIG S3E)**. Interestingly *BNI1* droplets also had the lowest circularity at 37°C but had the highest protein:RNA ratio at 25°C suggesting that the protein:RNA ratio is not a simple predictor of the material state in the system and suggests critical differences in molecular interactions must arise at high temperatures to alter material state. Notably, our previous work in other condensate forming systems (i.e. N protein of SARS CoV-2) shows that this association of viscoelastic properties at elevated temperature is not a general feature of protein-nucleic acid condensates but seems particular to the Whi3 system ^46^. Consistent with the more spherical shapes at low temperature, Fluorescence Recovery After Photobleaching (FRAP) on *CLN3* droplets at either 25°C or 37°C (**FIG S3H)** shows protein recovery was faster at 25°C and slower at 37°C, despite the higher thermal energy. Finally, we added fluorescently-labelled dextran molecules to *BNI1/* Whi3 condensates and measured whether the dextrans were enriched within the condensates or excluded from the condensates. We chose to use 10kDa dextran which has a hydrodynamic radius of around 3nm because previous work found that this sized dextran is enriched in reference Whi3 droplets at 30°C while larger dextrans (70kDa and 155kDa) were excluded^47^. We found that while dextran was enriched in droplets at 18°C and 25°C dextrans were excluded from condensates at 37°C indicating a distinct material state at high temperatures (**FIG S3I, J**). Thus, both isolates show temperature-dependent differences in the dense-phase concentrations of protein and RNA and differences in properties of droplets. These data support the idea that Whi3 protein and RNA condensates are temperature sensitive in a simplified *in vitro* system but as seen in vivo, the impact of temperature is not simply a binary shift in the C_sat_ for phase separation but is complex and impacts condensate composition and properties.

### Whi3 protein sequence is sufficient for a subset of temperature-sensitive activities

The *in vivo* and *in vitro* data suggest that Whi3 assemblies and the functions they control exhibit temperature-sensitive behavior. To assess if Whi3 protein sequence itself is sufficient to elicit temperature-sensitive phenotypes, we generated chimeric strains where we replaced the *WHI3* gene in one isolate with the *WHI3* sequence from different climate isolate to test if the Whi3 protein sequence was sufficient to rescue phenotypes seen at non-optimal growth temperatures. To create the chimeras, the full coding sequence of *WHI3* at the endogenous locus was exchanged between the warm and cold climate isolates generating a Florida isolate with a Wisconsin Whi3 (FL ^WI^ ^Whi3^) and a Wisconsin isolate with a Florida Whi3 (WI ^Fl^ ^Whi3^). In both cases Whi3 is tagged with an mNeon tag so that we could assess protein levels and observe condensate formation in the chimeric strains in addition to branching and nuclear synchrony phenotypes (**FIG S4A)**. We predicted that if key changes in Whi3 sequence were responsible for the temperature-dependent functions, the Whi3 replacement could be sufficient to rescue the temperature sensitivity of the “host” isolates.

We first assessed branching phenotypes in these geographic chimeras. In the WI isolate we found no major rescue of the branching phenotype at high temperatures when a FL Whi3 was present suggesting that a warm climate Whi3 was not sufficient to rescue the high temperature branching defect of cold climate-derived cells. Similarly, this exchange was insufficient to stimulate a low temperature sensitivity in the cold climate-derived cells (**FIG 5A**). Likewise, the WI-derived Whi3 was insufficient to rescue cold sensitivity in the FL isolates. The FL strain appeared to have some sensitivity to the Neon tag as some high temperature defects in branching were found that were not seen in untagged controls however these were relatively modest effects. Overall, we conclude that exchanging Whi3 sequences is not sufficient to rescue temperature sensitive phenotypes related to cell polarity.

We next tested whether exchanging Whi3 was sufficient to rescue temperature-sensitive mitotic synchrony defects. We measured the spindle pole body state in the two chimeric strains (Fl ^Wi^ ^Whi3^, WI ^FL^ ^Whi3^) and two control stains (Fl ^FL^ ^Whi3^, WI ^WI^ ^Whi3^) at 18°C, 25°C, 37°C. Both control strains performed similarly to the untagged strain with the WI isolate exhibiting synchrony defects when grown at high temperature and the FL isolate displaying synchrony defects at low temperature **(FIG 5B).** Remarkably, however, the cold climate isolate containing a warm climate Whi3 (WI ^Fl^ ^Whi3^) had a striking decrease in synchrony at high temperature, indicating a rescue of the phenotype. This strain also had a corresponding increase in synchrony at cold temperature, compared to the parent which had normal asynchrony in cold temperatures. Thus, the FL-derived Whi3 sequence was sufficient to both induce tolerance to the suboptimal warm temperature and also induce sensitivity to cold temperatures that the WI strain seemed adapted to tolerate. Similarly, replacing the FL Whi3 with the WI Whi3 (FL ^WI^ ^Whi3^) resulted in rescue of asynchrony in the cold and increase in mitotic synchrony at high temperature, thus mimicking the behavior of the WI-derived isolate rather than the warm-derived FL host. These replacement experiments show that the Whi3 protein sequence is sufficient to generate temperature-sensitive behavior in control of the nuclear division cycle (**FIG 5B**).

**Figure 5:**
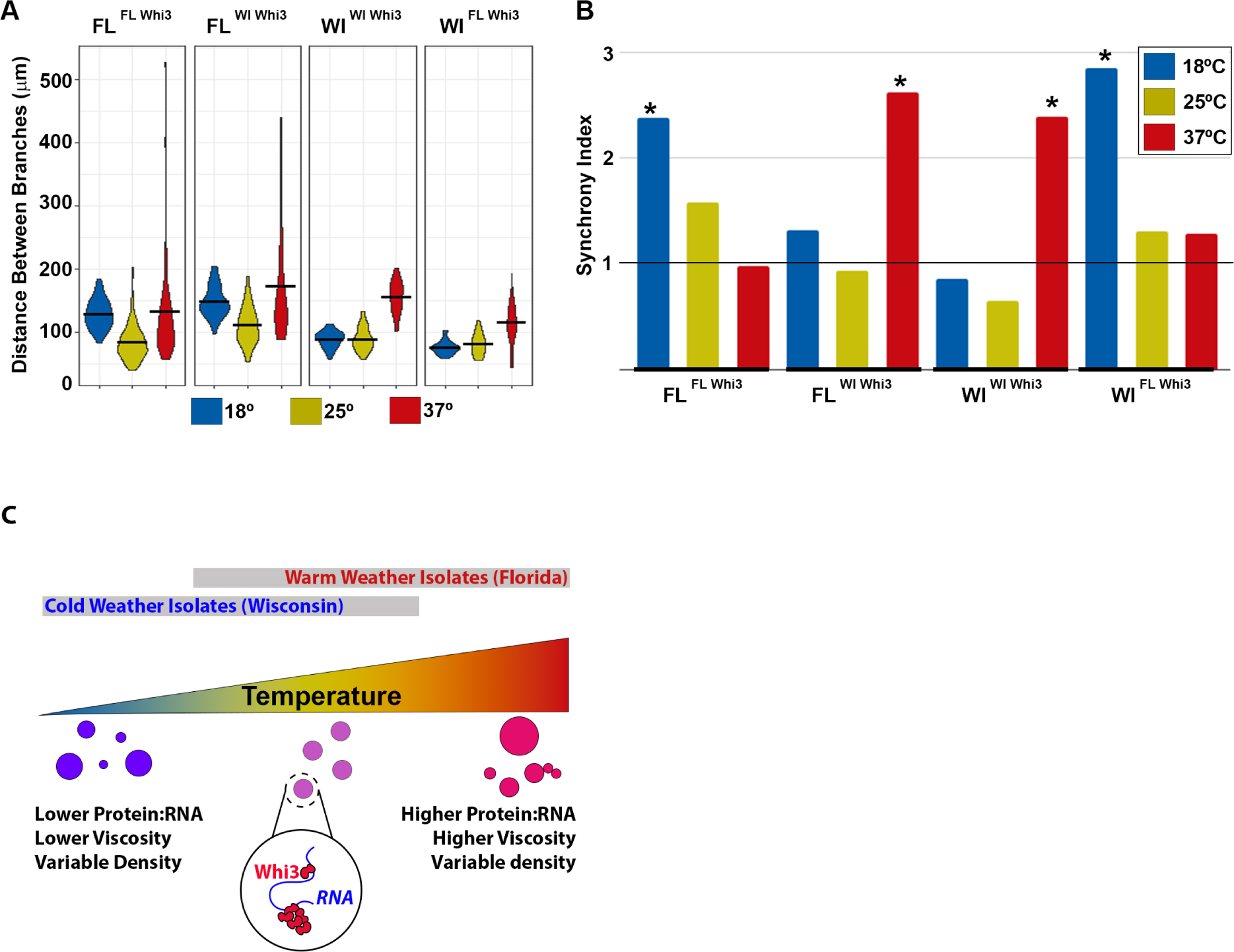
Whi3 protein is sufficient to rescue some temperature sensitive phenotypes **A)** Measurement of interbranch distance for each isolate and chimera at the indicated temperatures. **B)** Synchrony scores for each strain at different temperatures. Synchrony is calculated by dividing the proportion of nuclear pairs in which both nuclei are in mitosis divided the probability expected by change with 1 indicated random (t-test, * is p<0.05). **C)** Model for how temperature effect Whi3 condensates. Higher temperature leads to larger protein:RNA ration and more viscous droplets, lower temperature leads to more liquid droplets. Both conditions become less monodispersed. Due to sequence changes different Whi3 are still able to function despite these changes.

## Discussion

These results demonstrate that biomolecular condensates formed by Whi3 and RNA are temperature sensitive in their form and function. Our data suggests that the properties of these condensates are selected to function at specific temperatures and that a single component in the system can control the relationship between condensate function and temperature. Because Whi3 condensates in *Ashbya* are associated with several cellular processes it is possible to infer their functionality through clear phenotypic assays ^27,28^. We find that both branch initiation and nuclear cycle synchrony are altered in a temperature-dependent manner. Intriguingly, this includes loss of function phenotypes at both high and low temperature depending on the location of origin of the *Ashbya* strain. Notably, the protein Whi3 is sufficient to rescue a subset of temperature sensitive defects, despite the presence of many different polymorphisms between strains derived from different climates. This work shows that capacity of a small changes in an IDR to promote adaptation to growth in a new temperature regime ^27,28^.

A key observation in this study is that loss-of-function of condensates is associated with changes in their localization but not loss of higher-order assemblies. Despite the temperature-sensitive differences we see between isolates, we found no conditions in which condensates were completely absent *in vivo.* This contrasts with some other phase-separated compartments such as the *Drosophila* nucleolus or the *C. elegans* p-granule which can fail to form or dissolve at extreme temperatures ^11,12,14^. The presence of condensates suggests that the loss of function phenotypes are not due to the ability of the condensate to form but rather to the properties of the condensates that do form. A loss of function phenotype in the presence of condensates is consistent with other work in *Ashbya* in which loss of function phenotypes were created by mutating phosphorylation sites that never-the-less still had visible condensates in the proper locations^39^. These combined results suggest the material properties of condensates and whether they are functioning cannot be directly inferred from the simple, binary presence or absence of assemblies.

What then is the defect in these condensates at non-permissive temperature? How Whi3p functions differently in and out of condensates in *Ashbya* is not yet well understood. In both *Ashbya* and *S. cerevisiae,* RNA binding is critical for Whi3p function because mutations in the RRM domain phenocopy the null mutant. In *Ashbya* the formation of condensates is also critical because deletion of the QRR leads to severe phenotypes as well. Conversely, in yeast, deletion of the prion domain has no detectable phenotype on cell size in haploid cells^48^. Previous work in *Ashbya* has found that mutations that block the formation of condensates behave similarly to null mutants for branching and mitotic synchrony suggesting that the formation of condensates is essential for function. However, data from this study and elsewhere suggests that the presence of condensates is not sufficient for function^39^, and further, that larger condensates are associated with aberrant phenotypes. Data from yeast suggests that Whi3p can act as a translational repressor^49^. Unpublished data from our lab suggests Whi3 may promote or inhibit translation depending on conditions. We think these different roles in translation are associated with different scaled complexes, with potentially small condensates and/or soluble RNPs associated with promoting translation with larger condensates inhibiting translation.

The working functional model we favor is one in which Whi3 is responsible for positioning transcripts at certain locations in the cell. Whi3p then represses the translation of these RNAs. However, the condensates are likely under the control of cell cycle and polarity regulators that allow them to selectively release RNAs or dissolve completely at specific times for translation. This interplay allows the Whi3 condensates to position RNAs near tips and nuclei and compartmentalize the cytoplasm where those RNAs are eventually translated. When the temperature becomes non-permissive the properties of the droplets change and now we predict that these transient repressive interactions happen constitutively at a non-optimal time and place and result in the severe phenotypes at non-optimal temperatures. In this model, the function of the condensed state would be to impart timing control over the expression of resident RNAs with the key feature of the state being small scale assemblies that seem to be able to lead to transient repression. On-going work on spatial control of translation is testing this model.

While the Whi3 protein sequence was sufficient to rescue mitotic defects at non-optimal temperatures, it was insufficient to rescue polarity defects. Why is Whi3 not sufficient to rescue temperature-induced defects in polarity formation*?* One potential explanation is that while synchrony and polarity condensates both contain Whi3p they also contain other components. Condensates involved in polarity formation are known to contain the RNAs *BNI1* and *SPA2* and colocalize *in vivo* with Puf2p, in contrast condensates controlling synchrony are only known to contain Whi3p and *CLN3* ^27^. While this “parts list” may change as more research is done it is possible that Whi3 plays a larger role in determining the properties of nuclear-associated condensates than polarity condensates. Even with the known components, the target RNAs in polarity condensates are substantially longer than the *CLN3* RNA raising the possibility that the RNA sequence may be more critical in directing the temperature-sensitive function of these condensates.

One question raised by these results is whether there is a specific temperature module within Whi3. As seen in **FIG 2** most of the differences between isolates are located within the QRR. While our understanding of exactly how sequences of disordered domains function is incomplete, recent work in our lab has shed light on a potential mechanism. Adjacent to the QRR is a coiled-coil formed from 21 residues with leucines at the 1st, 8th, and 15th positions ^35^. This coiled-coil was found to strongly influence the material properties of droplets *in vitro* and was temperature sensitive *in vitro* moving from an alpha helical conformation to a disordered confirmation. Interestingly in *Ashbya* with an expanded QRR including the cold climate isolates there is a mutation in this domain that removes one of the leucines (**FIG S1B**) potentially causing this domain to behave less like a coiled-coil and more like a disordered region. Further studies will test these possibilities with targeted mutations.

Another non-mutually exclusive hypothesis is that the temperature-sensitivity is controlled by RNA in relation to the Whi3 protein. RNA structure is known to be temperature dependent ^50,51^, and RNA structure is known to affect the properties of droplets *in vitro* in the absence of sequences changes ^32^. Unpublished work from our lab found that *CLN3* molecules engineered to form more stable secondary structure result in condensates that are more liquid-like than *CLN3* molecules engineered to have less secondary structure. These changes in properties are similar to what we observe as temperature changes *in vitro* and could be one mechanism that leads to the counter-intuitive changes to material properties with temperature.

Free-living organisms cannot control their own temperature and must be able to adapt to a variety of fluctuations of varying magnitude that occur on the timescales of seconds to decades. How these changes affect the crucial functions performed by biomolecular condensates is not well understood but is likely critical to organism survival in many conditions. Overall, this work suggests that Whi3 condensates in the free-living fungus *Ashyba* are temperature sensitive and that the specific temperatures they are adapted to vary between isolates from different climatic regions. This study offers a potential mechanism by which free-living microbes can adapt growth to temperature through acquisition of relatively small numbers of mutations in central regulatory proteins. These results reveal the potential for rapid adaptation to fluctuating climate conditions via changes in IDRs of proteins that engage in condensates.

## Acknowledgements

This work was supported by NIH grant 7R01GM081506-13 to ASG, 1F32GM133123-01 to BMS, 1F32GM147989-01 to APJ. We would like to thank the Gladfelter lab for helpful discussions in the course of this work.

## Author Credit

Conceptualization: BMS, ASG

Funding Acquisition: BMS, ASG

Investigation: BMS

Methodology: APJ, SC, IS

Formal Analysis: FSD, GAM

Resources: LKF, FSD

Software: LKF, GAM

Visualization: BMS

Writing: BMS, ASG

## Methods

### Materials Availability

All unique stable reagents are available from the lead contact without restrictions.

### Branching and growth assay

*Ashbya* spores were plated onto Ashbya Full Media (AFM) for wild-isolates or AFM with Nourseothricin Sulfate (GoldBio, N-500-100, St. Louis. MO, USA) for wild-isolates with tagged Whi3. Spores were spread evenly across the plate surface using sterile glass beads and then plates were grown for 16-36 hours at either 18°C, 25°C, or 37°C. When spores had germinated they were photographed on a Zeiss Axioskop 40, using a Pixelink camera attached to a computer running PixelLink Capture software. Care was taken to ensure that each mycelium fit within a single frame, that each hyphae was distinguishable, and that each mycelium was the product of a single spore. Interbranch distance was measured using FIJI. Briefly, a freehand line was drawn from the tip of a hyphae to the first branch or between consecutive branches along a hyphae and the measure function was used to record the length of that line. Branch length was averaged per mycelium and at least one hundred cells were recorded per condition.

### Synchrony imaging and analysis

*Ashbya* spores were grown in liquid AFM for 16-32 hr in a baffled flask with shaking at 110 RPM at 18°C, 25°C, or 37°C. Cells were fixed with formaldehyde at a final concentration of 3.7% for 1hr, washed 3x with PBS and then resuspended in Solution A (100mK2PHO4, pH 7.5, 1.2M sorbitol) for digestion with zymolyse at 37°C to remove the cell wall. Cells were blocked for 1hr in PBS with 1mg/ml and 0.1% triton and then stained overnight with anti-alpha Tubulin antibody (1:100, MCA78G, BioRad). Cells were then washed, blocked again and stained with secondary antibody (Goat-Anti-Rat conjugated to Cy3 AP136C, SigmaAldrich) and DAPI (D9542, SigmaAldrich) and then mounted with vectashield (1551231, Fisher Scientific,).

Cells were imaged on a Nikon Ti-E stand equipped with a Yokogawa CSU-W1 spinning disk confocal unit, a Plan-Apochromat 100x/1.49 NA oil-immersion objective, and either a Prime 95b sCMOS camera (Photometrics) or a Zyla sCMOS camera (Andor). Nikon NIS-Elements software v.4.60. The spinning disk was illuminated with separate 405 and 561 nm laser sources. Images were scored by determining the cell cycle stage for each nucleus where possible based on the tubulin staining pattern. Because the staining pattern in mitosis is less ambiguous than in other cell cycle stages we focused on this stage of the cell cycle. We counted the total number of nuclei and the total number of nuclei in mitosis. We also counted all pairs of adjacent nuclei and counted the number of adjacent nuclei that were both in mitosis. We then calculated a synchrony score which was the actual number of paired mitotic nuclei divided by the number of pairs expected by chance. If the nuclei behave independently of each other this number will equal 1 with higher numbers indicating mitotic nuclei are more synchronous with their neighbors than expected by chance.

### *In vitro* RNA synthesis

Templates for *BNI1* RNA synthesis were generated by PCR from the genomic DNA with primers specific to the BNI1 gene that were compatible with all isolates (for: cgagtttttcagcaagattaatacgactcacgggacgacgacaagccta and rev: agatcttctagaaagatcctctctgatctggcgctaatgctc, IDT, Research Triangle Park, NC, USA). This fragment (∼6800bp) was gel purified using the QIAquick gel extraction kit according to the manufacture’s instructions (28706, Qiagen, Hilden, Germany) then cloned into a pJet vector using the clone jet kit (K1231 Thermo Fisher Scientific, Waltham, MA, USA) and then transformed into competent cells (C2987, New England BioLabs, Ipswich, MA, USA). Individual colonies were grown overnight under selection and DNA was extracted by miniprep according to the manufacturer’s instructions but without the presence of RNAse A in the resuspension buffer (FERK0503, Fisher Scientific, Hampton, NH, USA) Correct sequence was verified by Sanger sequencing (Genewiz, South Plainfield, NJ, USA). Plasmids were linearized using NcoI (R0197, R3193, New England BioLabs, Waltham, MA, USA) at 37°C for 1 hr. Linearization was confirmed by running the samples on an agarose gel. Linearized plasmids were purified using a QIAquick PCR purification kit according to the manufacturer’s instructions (28106, Qiagen, Hilden, Germany). Templates for *CLN3* RNA synthesis were in the same manner but using *CLN3* specific primers (for: TAATACGACTCACTATAGGGAGAGTCTGCATACCAAAGATCAGCCGCTTGC and rev: GTATGCTAGCGTAGTTTCTTGACC, IDT, Research Triangle Park, NC, USA). Linearized DNA for RNA transcription was generated by performing PCR on the pJET vector using the same plasmid and primers. Linearized DNA template was used to synthesize RNA according to the manufacturer’s instructions (M0658, New England BioLabs, Waltham, MA, USA). Briefly 1ug of DNA template was added to a tube with 2ul of reaction buffer, 2µl of Hi-T7 polymerase, 0.5ul of RNAse inhibitor, 1µl of NTP mix, 0.1µl of Cy-5 labeled UTP (PA55036, Sigma-Aldrich, St Louis, MO, USA) and enough water to bring the total volume to 20µl. RNA synthesis took place at 50°C for 1hr. The solution was diluted to 50ul and then the DNA was digested by addition of 2ul of DNAse (M0303, New England BioLabs, Waltham, MA). The RNA was precipitated by addition of 25µl of 2.5M LiCl and then chilled at -80°C for 30min. The RNA was pelleted by centrifugation at maximum speed for 10min, washed briefly with 70% Ethanol and then resuspended in nuclease free water. The concentration was determined by A260 measuring using a Nanodrop. Purity was assessed by denaturing gel electrophoresis.

### Protein synthesis and purification

Whi3 was cloned from the genomic DNA with primers specific to the Whi3 coding sequence (For: ctttaagaaggagatatacatatgcaccatcatcatcatcatatgtcgctggttaacagt, Rev: aagcttgtcgacggagctcgtcaagatttgccgaaggc IDT, Research Triangle Park, NC, USA). PCR products were run on an agarose gel and the band of the correct size (∼2200bp depending on isolate) was gel purified using the QIAquick gel extraction kit according to the manufacture’s instructions (28706, Qiagen, Hilden, Germany). The pet30-B vector was digested with Nde1 (R0111, New England BioLabs, Waltham, MA) and EcoRI-HF (R3101 New England BioLabs, Waltham, MA) and run on an agarose gel and a band of 5246bp was excised and gel purified. These fragments were combined using a the NEBuilder Hi-Fi DNA assembly kit according to the manufacture’s instructions (E5520, New England BioLabs, Waltham, MA). The resulting product was transformed into then transformed into competent cells (C2987, New England BioLabs, Ipswich, MA, USA). Individual colonies were grown overnight under selection and DNA was extracted by miniprep according to the manufacturer’s instructions (FERK0503, Fisher Scientific, Hampton, NH, USA) Correct sequence was verified by Sanger sequencing (Genewiz, South Plainfield, NJ, USA).

For protein synthesis the verified plasmid was transformed into BL21 competent cells (C2530, New England BioLabs, Ipswich, MA, USA). Cells were grown shaking at 37^°^C in 2XYT with appropriate selection to an OD600 of 0.6 and then transferred to 18^°^C. At this point cultures were induced with 0.1M of IPTG (I2481C25, GoldBio, St. Louis, MO, USA) and left to grow for 16hr. Cells were pelleted by centrifugation at 13000 rcf. The pellet was resuspended in 25ml Lysis buffer (1.5M KCl, 10mM Imidazole, 5mM beta-mercaptoethanol, 50mM Hepes pH7.4) along with 1 tablet Pierce Protease inhibitor (Thermo-FIsher Scientific A32965) and 15mg Lysozyme (Fisher BP535-10). Cells were lysed at 4°C with gentle rocking for 1hr and then briefly sonicated on ice. Lysate was clarified by centrifugation at 50000rcf for 30min. Clarified lysate was mixed with 2mL of Cobalt resin (Thermo Fisher Scientific PI89965), column was washed with 20 column volumes of lysis buffer and then eluted with 3ml of elution buffer (150mM KCl, 200mM Imidazole, 5mM beta-mercaptoethanol, 50mM Hepes pH7.4). Protein was dialyzed into storage buffer (150mM KCl, 50mM Hepes pH 7.4) using a slide-a-lyzer cassette and a 20,000MW cutoff (Thermo-Fisher Scientific 87735). Protein was labeled using an atto-488-NHS ester (Sigma Aldrich 41698) and then dialyzed again to remove unconjugated dye. Protein was concentrated just prior to use using a centrifugal filter (Amicon, UFC501024, Cork, Ireland). Protein concentration was determined using the Bradford Assay (Bio-Rad, 5000006, Hercules, California) according to the manufacturer’s instructions with measurements taken on a Nanodrop.

### *In vitro* condensate formation

For phase separation assays, glass-bottom imaging chambers (Grace Bio-Labs) were blocked with 30 mg/mL BSA (Sigma) in Whi3 buffer (150mM KCL, 50mM Hepes, 5mM beta-mercaptoethanol, pH 7.4) for 30 min to prevent protein adsorption to the surfaces of the well. The surfaces were washed thoroughly with Whi3 buffer prior to addition of protein and mRNA solutions. Whi3 buffer was added followed by Whi3 stocks followed by RNA to a final concentration of 1µM Whi3, 4 nM RNA and all wells were mixed at and then incubated at the indicated temperature. The final solution was mixed by pipetting without introducing bubbles. Imaging of *in vitro* condensates was performed with a Zeiss LSM 980 with Airyscan 2 using a Apochromat 63x 1.40NA Oil immersion objective. Illumination was done using a 488nm, and 639nm diode lasers. Microscope control and analysis was performed on premium HP Z6 workstations running Zeiss ZEN 3.7. To analyze images of condensates we built a custom FIJI macro. In brief, the macro thresholds each image based on the protein channel and then uses the built in “analyze particle” function to generate an ROI for each condensate. The macro then measures the size, shape, and intensity of each condensate. All condensates were generated in three separate wells and four images were taken from each well for quantification.

### Tagging of endogenous Whi3

To generate constructs to express fluorescent Whi3 at the endogenous locus and generate fluorescent Whi3 geographic chimeric strains, Whi3 was cloned from genomic DNA into a vector with an in-frame C-terminal mNeon tag, a NAT resistance gene, and homology to the the 3’ UTR of Whi3. Briefly: Whi3 and the 5’ UTR were amplified by PCR (for: attgggtaccgggcccccccgtttaaacagcatatttttcttttaattagattacacc rev: cacctgctccagatttgccgaaggcagc). mNeon was cloned from gDNA using primers (for: cggcaaatctggagcaggtgcaggtgca, rev: catacgtaatgctcaaccggttacttgtacagctcgtccatgc) the 3’ UTR of Whi3 was cloned from the genomic DNA (for: tgtttggctgcaggcatgcatgaaaatacgatatttgttcatttac rev: aaaagctggagctccaccgcgctcttcttaactatatatcttatcctttgcc). The backbone pRS416 was digested with SpeI, XhoI, and MfeI (R3133, R0146, R3589, New England BioLabs, Waltham, MA) generating fragments of 4841bp, 2536bp, 2073bp and 1746bp. The longest fragment was gel purified and saved. The NAT resistance gene was generated by digested pRS416 with AscI and HindIII (R0558, R0104, New England Biolabs, Waltham, MA) to generate fragments of 9467bp and 1729bp. The Shorter fragments was gel purified. Those five fragments (Whi3, mNeon, NAT, Whi3 3’UTR, Backbone) were combined using a the NEBuilder Hi-Fi DNA assembly kit according to the manufacturer’s instructions (E5520, New England BioLabs, Waltham, MA). The resulting product was transformed into then transformed into competent cells (C2987, New England BioLabs, Ipswich, MA, USA). Individual colonies were grown overnight under selection and DNA was extracted by miniprep according to the manufacturer’s instructions (FERK0503, Fisher Scientific, Hampton, NH, USA) Correct sequence was verified by Sanger sequencing (Genewiz, South Plainfield, NJ, USA) and PacBio Sequencing (Plasmidsaurus, Eugene, OR, USA).

The resulting plasmids were mini-prepped and then digested with PmeI and SapI (R0560, R0569, NewEngland BioLabs, Waltham, MA) to generate a 6.01kb fragment containing the Whi3 coding sequence and 5’ UTR, mNeon, and Nourseothricin sulfate resistance, and the Whi3 3’UTR. This fragment was transformed into *Ashbya* by electroporation as previously described (Wendland et al., 2000). Transformants were verified by PCR and Sanger sequencing (Genewiz, South Plainfield, NJ, USA).

For imaging, cells were grown overnight on AFM agar plates containing Nourseothricin Sulfate at 18°C, 25°C, or 37°C. Cells were then examined under a dissecting scope and suitable colonies were identified. Using a razor blade a 5mm x 5mm square of agar containing these colonies was cut from the plate which was then inverted onto a coverslip. Cells were imaged on on a Nikon Ti-E stand equipped with a Yokogawa CSU-W1 spinning disk confocal unit, a Plan-Apochromat 100x/1.49 NA oil-immersion objective, and a Zyla sCMOS camera (Andor). The microscope was equipped with a temperature-controlled microscope stage (Tokai Hit from Incubation System for Microscopes) to control the temperature while imaging. Nikon NIS-Elements software v.4.60 was used to control the machine. The spinning disk was illuminated with a 488 nm laser source.

### Data Analysis

All data analysis was performed in R Studio with R 3.3.0.

