## Supplemental Figures for "Biomolecular condensates in fungi are tuned to function at specific temperatures"

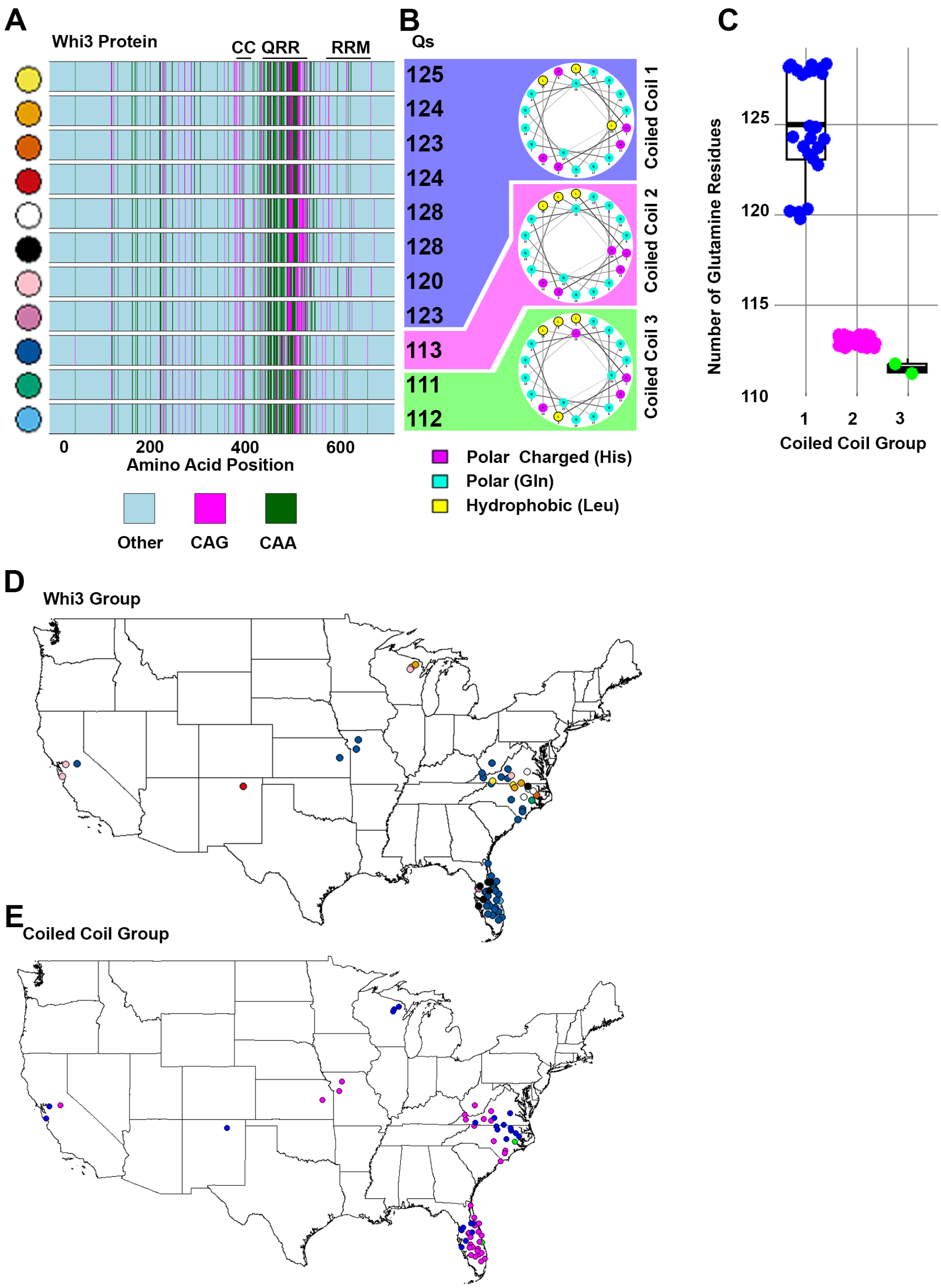


**Figure S1: A)** Complete sequence alignment for all unique Whi3 sequences showing Glutamines in magenta (for CAG codon) and green (for CAA codon) plus all other residues (blue). The total number of glutamines in the sequence is noted to the right. **B)** Helical diagrams showing the three varieties of coiled coil domain identified in Seim *et al.* 2022. Colors on the wheel represent polar but uncharged glutamine (cyan), polar positively charged histidine (purple), and nonpolar hydrophobic leucine (yellow). Hydrophobic residues cluster on a single face in coiled variety 2 and 3 but are interrupted with a polar residue in coiled coil variety 1 **C)** Boxplots comparing the number of glutamines in the Whi3 sequence compared to the coiled coil class from B. Variety 2 and 3 containing an intact coiled coil and have fewer total glutamines. (**D)** Map of the locations of each isolate. Colors correspond to the left side of panel A. **E)** Location of the different coiled coil varieties show in panel B.

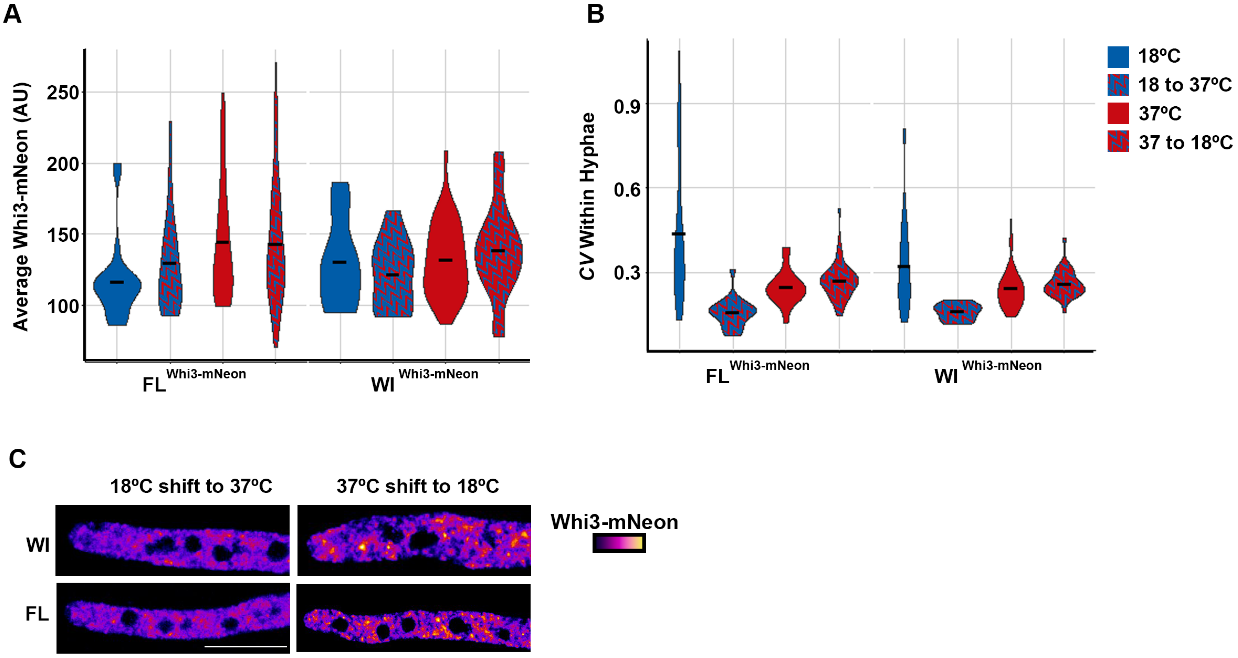


**Figure S2**: Temperature shifts affect protein localization but not total protein abundance in cells. **A)** Average Whi3-mNeon signal per hyphae when grown at 18°C (blue), grown at 18°C and shifted to 37°C (blue with red stripes), grown at 37°C (red), or grown at 37°C and shifted to 18°C (red with blue stripes) for each isolate. Protein concentration within hyphae did not change following shifts. **B)** CV of Whi3-mNeon signal per hyphae. **C)** Representative images of hyphae from the indicated location grown at 18°C and shifted to 37°C or grown at 37°C and shifted to 18°C. Scale bar represents 10µm.

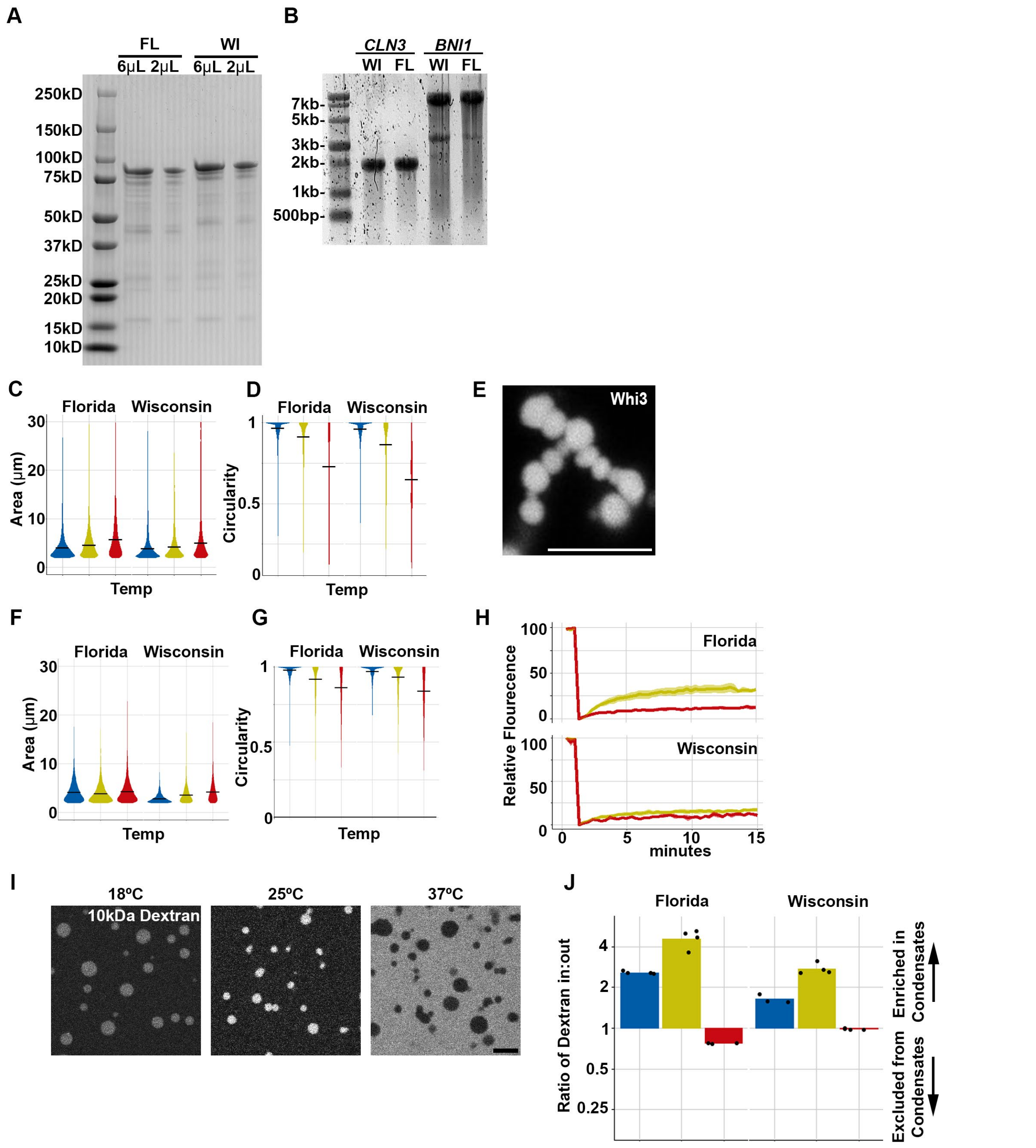


**Figure S3: A**). Coomassie Blue stained gel showing purified Whi3 protein from Florida and Wisconsin isolates. **B**) Denaturing RNA gel showing *in vitro* synthesized RNA of the indicated genes from Wisconsin and Florida. **C**) Area of each Whi3+*CLN3* droplet from the indicated strain and temperature. **D)** circularity calculated as 4*π (area / perimeter$^2$) for each droplet at the indicated strain and temperature. **E**) Close up of Whi3/*BNI1* condensate at 37°C showing numerous small droplets fused together but not relaxed. **F**) Area of each Whi3/*BNI1* droplet from the indicated strain and temperature. **G**) Circularity calculated as 4*π (area / perimeter$^2$) for each droplet at the indicated strain and temperature. **H)** Fluorescence recovery after photobleaching (FRAP) curves for Whi3/*BNI1* condensates at the indicated temperature and from the indicated strains. **I**) Rhodamine-labelled dextran of molecular weight 10kDa were added to

Whi3/*BNI1* condensates at each indicated temperature. **J**) Ratio of dextran fluorescence within condensates and outside of condensates from each isolate at the indicated temperature. The location of condensates was determined by observing atto-488 labeled Whi3 (not shown). A ratio of 1 indicates homogenous signal, greater than 1 indicates dextran was condensate enriched, a value less than 1 indicates dextrans are condensate excluded. scale bars are 5μm.

**A**

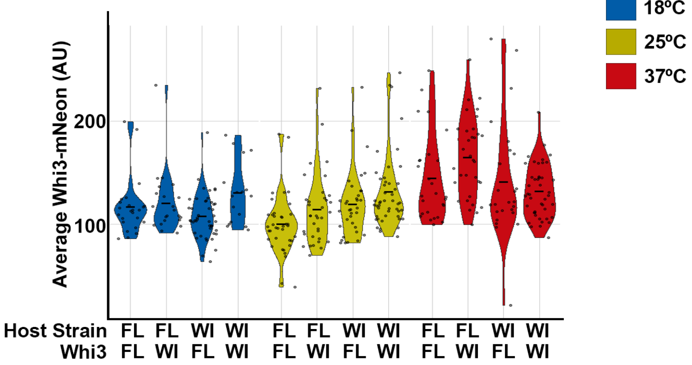


**Figure S4: A**) Average Whi3-mNeon protein level per hyphae at the indicated temperatures.
